# CD24a knockout transforms the tumor microenvironment from cold to hot by promoting tumor-killing immune cell infiltration in a murine triple-negative breast cancer model

**DOI:** 10.1101/2024.07.15.603489

**Authors:** Shih-Hsuan Chan, Hsuan-Jung Tseng, Lu-Hai Wang

## Abstract

**Background:** CD24 plays a crucial role not only in promoting tumor progression and metastasis but also in modulating macrophage-mediated anti-tumor immunity. However, the impact of tumor CD24 on the immune landscape of the tumor microenvironment (TME) remains poorly explored. Here, we investigated the role of CD24a, murine CD24 gene, in the progression and immune dynamics of the tumor microenvironment (TME) in the 4T1 murine model of triple-negative breast cancer (TNBC). **Methods:** We employed Clustered Regularly Interspaced Short Palindromic Repeat (CRISPR)/Cas9 technology to perform a gene knockout of Cd24a in 4T1 cells. Flow cytometry was utilized to analyze the distribution and number of immune cells, including myeloid-derived suppressor cells (MDSCs), natural killer (NK) cells, T cells, and macrophages, within tumors, spleens, and bone marrow. Immunofluorescence (IF) staining was used to detect these immune cells in tumor sections. Additionally, ANDOR Dragonfly High-Speed Confocal was used to perform three-dimensional (3D) mapping of mouse tumors.

**Results:** Our study showed that knocking out CD24a significantly impeded tumor progression and prolonged mouse survival. Flow cytometry and IF analysis revealed that the loss of CD24a transformed tumor microenvironment from cold to hot by promoting the infiltration of M1 macrophages, cytotoxic CD8^+^ T cells, and CD49b^+^ natural killer (NK) cells while reducing the recruitment and expansion of granulocytic myeloid-derived suppressor cells (gMDSCs) in the TME. Additionally, the 3D mapping of TME further validated the “hot state” of CD24a knockout tumors.

**Conclusions:** Our study provides the first evidence that targeting CD24a could effectively reprogram the TME, enhancing its immunogenicity, and transforming immune cold tumors into hot tumors. This strategy may offer a promising therapeutic strategy for enhancing the immune response against poorly immunogenic tumors.

## Introduction

The CD24 gene encodes a highly glycosylated glycosylphosphatidylinositol (GPI) anchored membrane without a transmembrane domain [1, 2]. CD24 has been shown to play diverse functional roles in immunity, cancer, inflammation, and autoimmune diseases [2]. CD24 is predominantly expressed on B-cell progenitors, with a lesser presence on terminally differentiated B cells. It functions as a costimulatory molecule on activated B cells, facilitating the clonal expansion of CD4 T cells [3–5]. Furthermore, CD24 was identified as an immune modulator to suppress in the danger-associated molecular patterns (DAMPs)-induced inflammatory signals, which could be lethal during host infection [6]. Prior investigations have established that the interaction between CD24 and Siglec-10 has the capacity to suppress DAMP-mediated inflammatory responses, contributing to the immunosuppression observed in placental tissues [6]. In addition, CD24 overexpression has been associated with several hallmarks of cancer, including increased cell proliferation, migration, and metastasis [3, 7]. CD24’s interaction with P-selectin has been demonstrated to facilitate cancer cell adhesion to endothelial cells, promoting pulmonary metastasis [8, 9]. Additionally, CD24 serves as an oncogenic signaling activator, influencing pathways such as lipid raft/integrin-induced Src signaling [10], and the Wnt/β-catenin pathway [11], and the EGFR/MET/Akt/Erk1/2 pathway [7, 12]. Our previous work on triple-negative breast cancer (TNBC) revealed the involvement of CD24 in TNBC progression and metastatic lung colonization, mediating anoikis resistance and angiogenesis through RTK oncogenic pathways [12]. On the other hand, a number of studies have shown that CD24 is an unfavorable marker for ALDH1-positive CD44^+^/CD24^-^ breast cancer initiating cells (BCICs) [2, 13, 14]. Aside from breast cancer, several studies have identified CD24 as an ovarian cancer stem cell marker, which is pivotal in ovarian cancer progression and metastasis [15, 16].

In addition to CD24’role in cancer progression, recent research has shed light on CD24’s function as a “don’t eat me” signaling molecule that interacts with macrophage Siglec-10, triggering a signaling cascade that inhibits macrophage-mediated phagocytosis [17]. This “don’t eat me” signal is crucial for tumor cells to evade macrophage-mediated anti-tumor immunity, potentially contributing to the creation of an immunosuppressive tumor microenvironment (TME). However, whether the presence of CD24 directly contributes to the formation of an immunosuppressive TME remains to be elucidated. In this study, we employed the CRISPR/Cas9 genome-editing technique to knockout the *Cd24a* gene in 4T1 cells, a murine model of TNBC, and this allowed us to investigate the effects of CD24a loss on the potential alteration of immune landscape of TME in an orthotopic breast cancer mouse model. Our goal was to elucidate how CD24a loss influences immune landscape of TME and the subsequent tumor progression.

## Materials and Methods

### Cell lines and reagents

Murine breast cancer cell line 4T1 was cultured in 10% fetal bovine serum (FBS) Dulbecco’s Modified Eagle Medium (DMEM) (Life Technology, CA, USA) and was incubated in a humidified incubator at 37℃ supplied with 5% CO^2^. OPTI-MEM medium was purchased from Life Technology Inc. Mirus LT1 transfection reagent was purchased from Mirus company (Life Technology, CA, USA). Culture supernatant was sampled and examined for mycoplasma contamination every one month. Cell identity was verified using STR analysis. Reagents and antibodies used in this study were listed in the supplementary table S1.

### Knockout (KO) of *Cd24a* gene in 4T1 cells using Clustered Regularly Interspaced Short Palindromic Repeat (CRISPR)/Cas9 technology

Two sgRNAs targeting the upstream of ATG start codon and downstream of exon 1 of *Cd24* gene on mouse genomic DNA were synthesized and cloned into All-in-one system sgRNA/Cas9 expression lentivector using BsmbI restriction enzyme site. The sequences of sgRNAs were as follow: *Cd24a* sgRNA-1: AGAGTCGCGCCGCGCGCCGA and *Cd24a* sgRNA-2: GGCACTGCTCCTACCCACGC. Two sgRNA/Cas9 expression lentivectors were transiently co-transfected into 4T1 cells using Mirus® XT transfection reagent for eight hours followed by puromycin selection to remove the untransfected cells. The surviving colonies were subjected to a single cell culture and the individual clone grown from a single cell culture was screened for the loss expression of CD24 using flow cytometry. Clones whose CD24 expression were completely lost were subjected to the extraction of genomic DNA. We designed a primer pair to PCR amplify the genomic sequences of 804 bp containing 200 bp upstream of the first sgRNA targeting site and 200bp downstream of the second sgRNA targeting site. DNA sequencing of the PCR product of each positive clone was carried out using Sanger sequencing method.

### Flow cytometry analysis

To isolate immune cells from tumors, the resected tumors were chopped into pieces and subjected to enzyme digestion with a solution containing Accutase and 0.25% trypsin in a 1:1 ratio for one hour at 37°C. The digested tumor pieces were passed through a cell strainer, and the tumor-infiltrating cells were further isolated using density gradient centrifugation with Ficoll-Paque™ PLUS (Cytiva, CA, USA). The tumor-infiltrating immune cells were collected from the interface between PBS and Ficoll. For immune cell isolation from mouse spleens, the spleens were placed on a cell strainer set on a 50 mL tube and thoroughly crushed with a plunger, followed by washing with 5 mL of 1X phosphate-buffered saline (PBS) to isolate the single-cell population. The tube was centrifuged at 3,000 rpm for 5 minutes, the supernatant was discarded, and the cell pellet was resuspended with RBC lysis buffer. This was followed by another centrifugation at 3,000 rpm for 5 minutes to obtain the immune cells. To isolate mouse bone marrow cells, the hind long bones (femora) were dissected, and the ends were cut off with scissors. The bones were flushed with 2 mL of 1X PBS to collect the bone marrow cells, followed by RBC lysis buffer treatment. This was followed by another centrifugation at 3,000 rpm for 5 minutes to obtain the immune cells.

The populations of CD11b^+^Ly6C^+^ monocytic MDSCs (mMDSCs) and CD11b^+^Ly6G^+^ granulocytic MDSCs (gMDSCs) in tumor-infiltrating immune cells, splenocytes, and bone marrow cells were analyzed by staining the cells with FITC-conjugated anti-CD11b antibody, PE-Cy7-conjugated anti-Ly6C antibody, and PE-conjugated anti-Ly6G antibody at a 1:100 dilution. The population of CD11b^+^F4/80^+^ tumor-infiltrating macrophages in tumors was stained with FITC-conjugated anti-CD11b antibody and PerCP-Cy5.5-conjugated F4/80 antibody at a 1:100 dilution. The population of CD3^+^CD8^+^ tumor-infiltrating T cells in tumors and spleens was analyzing by staining with FITC anti-CD3 antibody, and Per/CP cy5.5-conjuated anti-CD8 antibody at a 100X dilution. The population of CD49^+^ tumor-infiltrating natural killer cells in tumors and spleens was stained with FITC-conjugated anti-CD49b^+^ antibody at a 100X dilution. Data acquisition was performed using BD Canto flow cytometry, and Flow Jo software was utilized for subsequent data analysis.

### Tumor spheroid formation assay

A total of 1 x10^5^ cells were resuspended and seeded in the ultra-low 6-well plates with MACSsphere medium. The cells were cultured for 7 days in a humidified incubator at 37℃ supplied with 5% CO2. The images of tumor spheroids were photographed using light microscopy and were calculated using Image J software.

### MTS cell proliferation assay

3 x 10^3^ 4T1 cells and CD24a KO cells were seeded in a 96-well plate respectively 24 hours prior to the experiment. The viability of cells was then evaluated using an MTS assay. The MTS reagent was introduced into the well and cells were incubated for 1 hour. Subsequently, the optical density was measured at 450 nm.

### Animal study

2 x 10^4^ parental 4T1 and *Cd24a* KO 4T1 cells were mixed with Matrigel in 1X PBS at a ratio of 2:1. These cells were then injected into the 4^th^ fat pad of BALB/c mice. Tumor growth was monitored weekly over a one-month period. After one month, the mice were sacrificed, and the tumors were excised. The resected tumors were fixed in 2% paraformaldehyde, embedded in optimal cutting temperature (OCT) compound on dry ice, and stored at −80℃ for subsequent immunofluorescence staining (IF) analysis. For the survival study, the parental 4T1 and *Cd24a* KO 4T1 cells were similarly injected into 4^th^ fat pad of BALB/c mice. The survival of the tumor-bearing mice was monitored over a two-month follow-up period.

### Immunofluorescence staining

The OCT-embedded mouse tumors were sectioned into 10 μm tissue slices using a freezing microtome. Briefly, the tumor slices were washed with 1X PBS three times to removed OCT and then were blocked in the blocking buffer (1X PBS containing 5% goat serum and 1% bovine serum albumin) for one hour at room temperature. The tumor sections were washed with 1X PBS to remove the blocking buffer followed by incubation of the specific primary antibodies overnight at 4℃. The tumor sections were washed with 1X PBS three times to remove the primary antibodies followed by incubation of the fluorescence-tagged secondary antibodies for one hour at room temperature. The tumor slices were washed with 1X PBS buffer and mounted with anti-fade mounting medium containing DAPI (Life technology Inc. CA. USA).

### Three-Dimensional (3D) imaging

Mouse tumors were fixed in 4% paraformaldehyde and sectioned into 500 μm slices using a 5100mz vibrating microtome (Campden Instruments Ltd., UK). Following fixation, the sections were washed three times with 1X PBS. The tumor sections were then permeabilized with 2% Triton X-100 overnight, followed by three additional PBS washes to remove Triton-X100. The tumor slices were blocked overnight in a blocking buffer containing 10% goat serum in 1X PBS. Primary antibodies were prepared in a 1:100 dilution in an antibody dilution buffer (1X PBS containing 1% goat serum and 0.25% Triton-X100) and incubated with the tumor sections for 48 hours at 4℃. Afterward, the sections were washed three times by a washing buffer (0.1% Triton-X100 1X PBS) at 4℃. The tumor slices were then incubated with the fluorescence-conjugated secondary antibodies (1:300 dilution in the antibody dilution buffer) for 24 hours at 4℃ followed by three PBS washes in a washing buffer (0.1% Triton-X100 1X PBS) at 4℃. Next, the tumor slices were stained with Hoechst (1:500 dilution) for 15 minutes followed by PBS wash three times at room temperature to remove Triton-X100. Finally, the tumor sections were treated with RapiClear CS solution (SunJin Lab Co., CA, USA) at room temperature for one hour to achieve the tissue clearing. The 3D image of the tumor sections was obtained using the ANDOR Dragonfly High Speed Confocal (Oxford Instruments, UK), and the images and video were analyzed and generated using Imaris 9.7 image analysis software.

### Statistical analysis

The student’s t-test was employed to examine differences between the means of two data sets. To compare the Kaplan-Meier survival curves of two groups, the Log-rank test was utilized. Furthermore, the Cox proportional hazards model was employed to estimate the hazard ratio between the groups. The statistical significance was defined as a p-value less than 0.05.

## Results

### CRISPR/Cas9 knockout of *Cd24a* gene in 4T1 murine breast cancer cells

To establish *Cd24a* knockout (ΔCD24a) 4T1 cells via the CRISPR/Cas9 method, we designed and inserted two sgRNAs downstream of the U6 promoter in Cas9-expressing plasmids. Specifically, one sgRNA targeted sequences upstream of the transcription start site (TSS) of the *Cd24a* gene, while the other targeted the exon 1-intron 1 junction of the *Cd24a* gene in the mouse genome. 4T1 cells were transfected with sgRNA-containing Cas9-expressing plasmids followed by puromycin selection. The *Cd24a* gene knockout status of the resulting stable cells was validated by Sanger sequencing, which revealed a successful deletion of exon 1 (Fig.1A) and confirmed by RNA sequencing (Fig. 1B). Flow cytometry was used to further validate the loss of membrane CD24a expression in two ΔCD24a 4T1 clones, ΔCD24a-1 and ΔCD24a-2, as compared to WT 4T1 cells (Supplementary Fig. S1).

**Fig 1.**
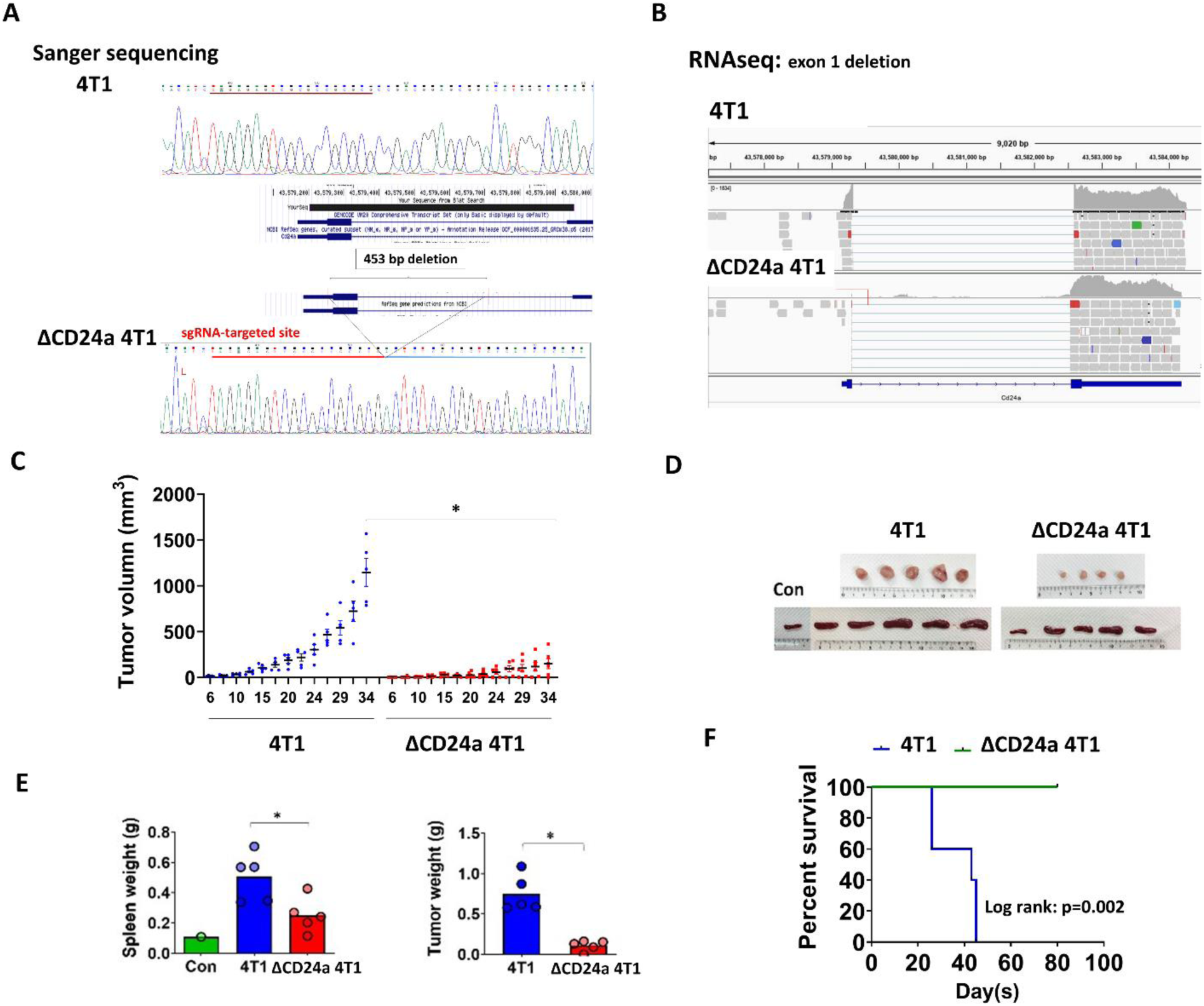
Knockout of CD24a significantly impairs tumor growth and prolongs the mouse survival in a 4T1 BALB/c syngeneic model. **A**, Sanger sequencing of _ΔCD24_ 4T1 cells revealed a 453 bp deletion containing exon 1 and half of intron 1 of *Cd24a* gene. **B**, RNAseq analysis showed that the coding region of *Cd24a* exon 1 was deleted in ΔCD24a cells as compared to the wild type 4T1 cells. **C**, The growth curve between WT (n=5) and ΔCD24a 4T1 tumor **(n=5)** in a Balb/c mouse model. **D**, Kaplan-Meier survival analysis of ΔCD24a 4T1 tumor-bearing mice and WT 4T1 tumor-bearing mice. **E**, Tumor growth kinetics of of ΔCD24a 4T1 tumor-bearing mice and WT 4T1-bearing mice. *P<0.01. **F**, The weight of tumors and spleens are shown in histograms. *P<0.01.

### CD24a Knockout impaired *in vitro* spheroid-forming ability and *in vivo* tumorigenicity of 4T1 cells

We first examined the effect of CD24a knockout on tumorigenicity of 4T1 cells. The spheroid formation assay was performed to evaluate the *in vitro* tumorigenicity of 4T1 cells. We found that CD24a knockout significantly impaired spheroid-forming ability of 4T1 cells (Supplementary Fig. S2A) without affect cell growth (Supplementary Fig. S2B). Subsequently, ΔCD24a-1 clone was selected for the following mouse study. In the 4T1/BALB/c syngeneic mouse model, we demonstrated that ΔCD24a 4T1 cells grew smaller tumors as compared with WT 4T1 cells (Fig. 1C). In addition, ΔCD24a 4T1 tumor-bearing mice showed significantly less splenomegaly than WT tumor-bearing mice (Fig. 1D and Fig. 1E). Moreover, ΔCD24a 4T1 tumor-bearing mice lived significantly longer than WT tumor-bearing mice (Fig. 1F p=0.002, HR: 0.049. 95% CI: 0.007-0.348), and the median survival was not reached. These data indicated that loss of CD24a significantly impaired tumor progression in the 4T1/BALB/c syngeneic mouse model.

### CD24a knockout significantly suppresses the recruitment of granulocytic MDSCs (gMDSCs) to the tumor microenvironment

Next, we examined the impact of CD24a loss on the immune landscape of TME. We first looked at the possible alteration of MDSCs population in the tumor, spleen, and bone morrow, in which they were produced. MDSCs are heterogenous immature myeloid cells known to accumulate in the TME during tumor progression and considered to be a strong contributor to the immunosuppressive tumor microenvironment [18, 19].

Tumors resected from mice were chopped with the surgical knife and minced with the plunger followed by accutase digestion at 37℃ for one hour. The digested cell mixtures were passed through the cell strainer to obtain single cell population. The isolated immune cells were then stained with antibodies specific for CD11b^+^Ly6G^+^ granulocytic MDSCs (gMDSCs) and CD11b^+^Ly6C^+^ monocytic MDSCs (mMDSCs). Flow cytometry analysis showed that ΔCD24a 4T1 tumor-bearing mice had decreased number of CD11b^+^Ly6G^+^ granulocytic MDSCs (gMDSCs) in the spleen and tumor but had similar number of gMDSCs in bone marrow as compared with WT tumor-bearing mice (Fig. 2A and 2B), suggesting that ΔCD24a 4T1 tumors were less potent to recruit gMDSCs from the bone marrow to the spleen and to the primary tumor site as compared with WT tumors. Meanwhile, ΔCD24a 4T1 tumor-bearing mice also showed decreased number of CD11b^+^Ly6C^+^ myeloid MDSCs (mMDSCs) in the spleen but had similar numbers of mMDSCs in the tumor and bone marrow (Fig. 2A and 2C).

**Fig. 2.**
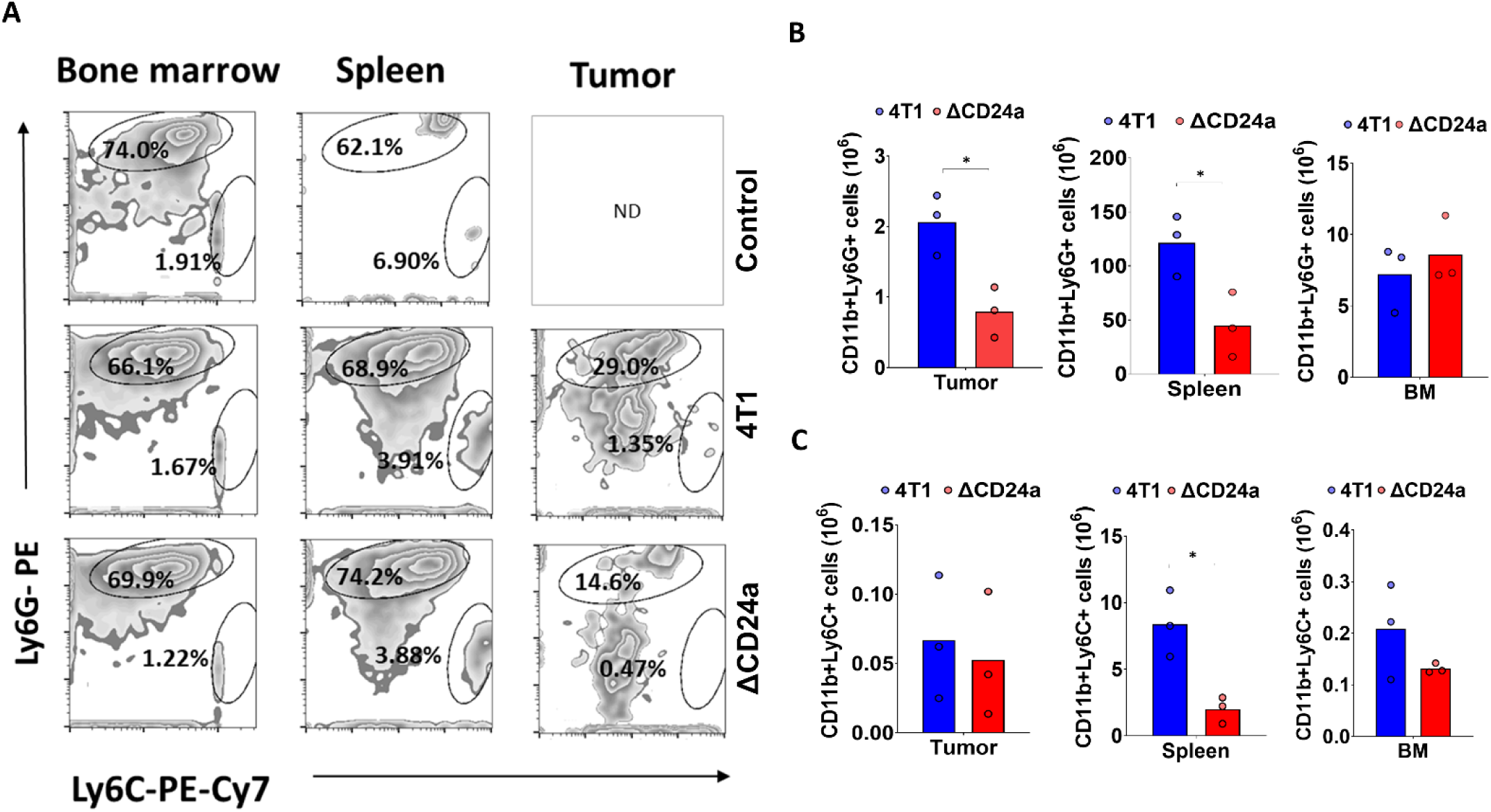
Knocking out CD24a notably reduces the generation of granulocytic and monocytic myeloid-derived suppressor cells (MDSCs) in the spleens and decreases the recruitment of gMDSCs to the tumor microenvironment. **A**. Flow cytometry analysis of the distribution of CD11^+^Ly6G^+^ granulocytic MDSCs (gMDSCs), and CD11b^+^Ly6C^+^ monocytic MDSCs (mMDSCs) (n=3) in the bone marrows, spleens, and tumors. **B**, The cell numbers of gMDSCs, and **C**, mMDSCs in the tumors, spleens and bone marrows are shown in histograms. **C**, The cell numbers of mMDSCs in the tumors, spleens and bone marrows are shown in histograms.

### CD24a knockout induces macrophage infiltration in and natural killer (NK) cells recruitment to the tumor microenvironment

Next, we investigated whether knocking out CD24a influences the infiltration of tumor-associated macrophages (TAMs), which are crucial in tumor progression, within the tumor microenvironment (TME). Using flow cytometry, we analyzed the presence of CD11b^+^/F4/80^+^ TAMs in ΔCD24a 4T1 tumors and WT tumors. The results indicated a significant increase in CD11b^+^/F4/80^+^ macrophage infiltration in the ΔCD24a 4T1 tumors compared to the WT tumors, showing a 2.5-fold increase in the number of tumor-infiltrating macrophages (Fig. 3A). Subsequently, we examined the distribution of natural killer (NK) cells in the spleens and tumors of mice. Flow cytometry analysis showed that ΔCD24a 4T1 tumors-bearing mice had lower numbers of CD49b^+^ NK cells in the spleen but higher cell densities inside the tumors as compared with WT tumor-bearing mice (Fig. 3B). These results suggested that knocking out CD24a in the tumor cells enhanced macrophage infiltration and NK cells recruitment to the tumor area.

**Fig. 3.**
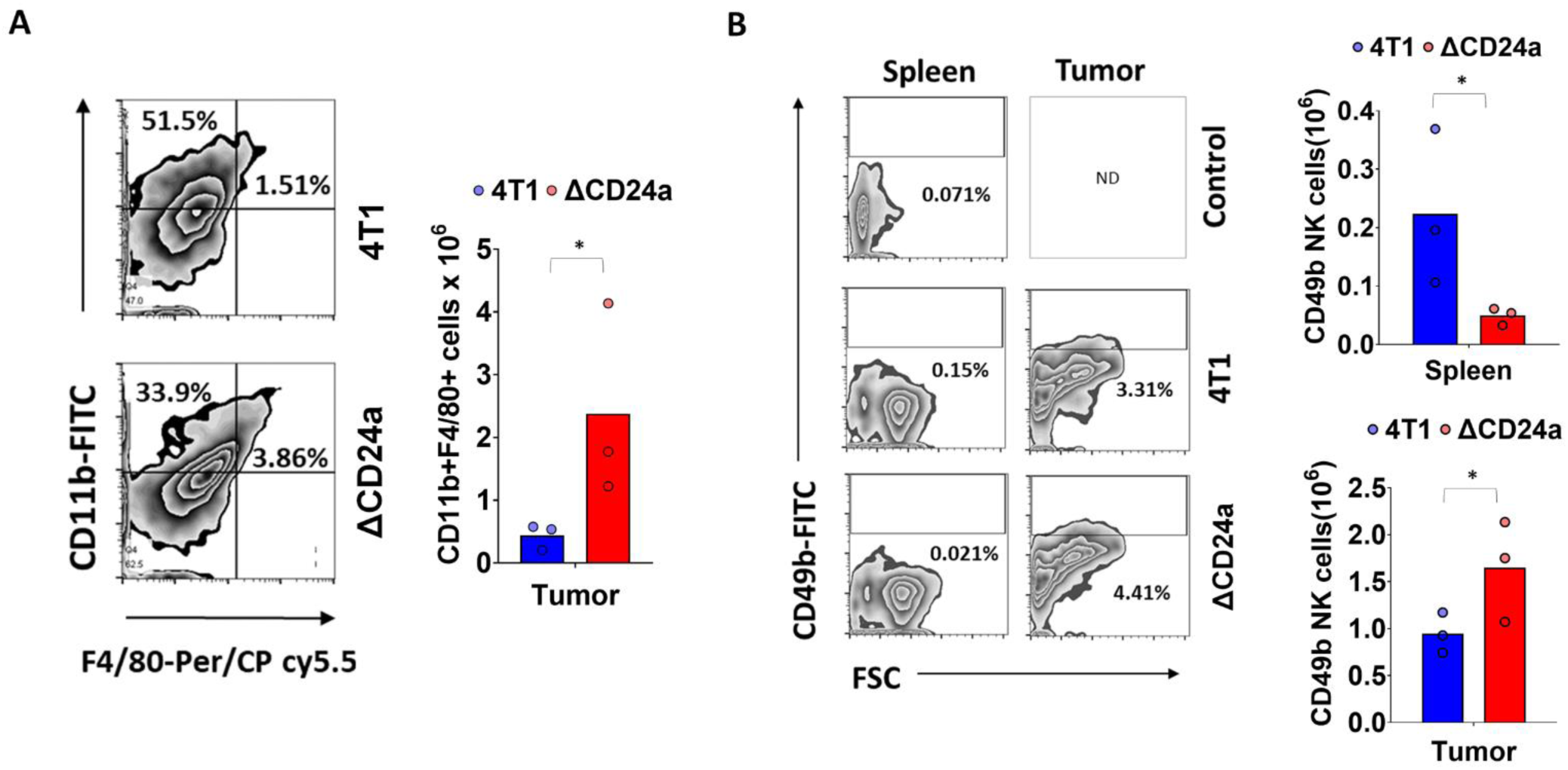
The loss of CD24a significantly enhances macrophage infiltration and the recruitment of natural killer (NK) cells to the tumor microenvironment. **A**. Flow cytometry analysis of tumor-infiltrating CD11b^+^ F4/80^+^ macrophages (n=3). The number of CD11b^+^ F4/80^+^ macrophages cells in the tumor and spleen is presented in histograms. *P<0.05. **B.** Flow cytometry analysis of the presence of CD49b^+^ NK cells in the tumor and spleen. The number of CD49b^+^ NK cells in the tumor and spleen is presented in histograms. *P<0.05.

### CD24a knockout significantly suppresses the infiltration of cytotoxic CD8^+^ T cells in tumor microenvironment

The reduced tumor burden observed in ΔCD24a tumor-bearing mice compared to WT tumor-bearing mice led us to ask how CD24a knockout affects cytotoxic CD8^+^ T cell infiltration, which plays a major role in tumor killing, in the tumor microenvironment. As depicted in the Fig. 4A and 4B, ΔCD24a tumors exhibited a significantly higher number of infiltrating CD8^+^ T cells compared to WT tumors. Conversely, ΔCD24a 4T1 tumor-bearing mice demonstrated lower densities of CD8^+^ T cells in the spleen compared to their WT counterparts (Fig. 4A and 4B). These findings demonstrated that the knockout of CD24a in tumor cells significantly enhances the recruitment and infiltration of cytotoxic CD8^+^ T cells at the tumor site.

**Fig. 4.**
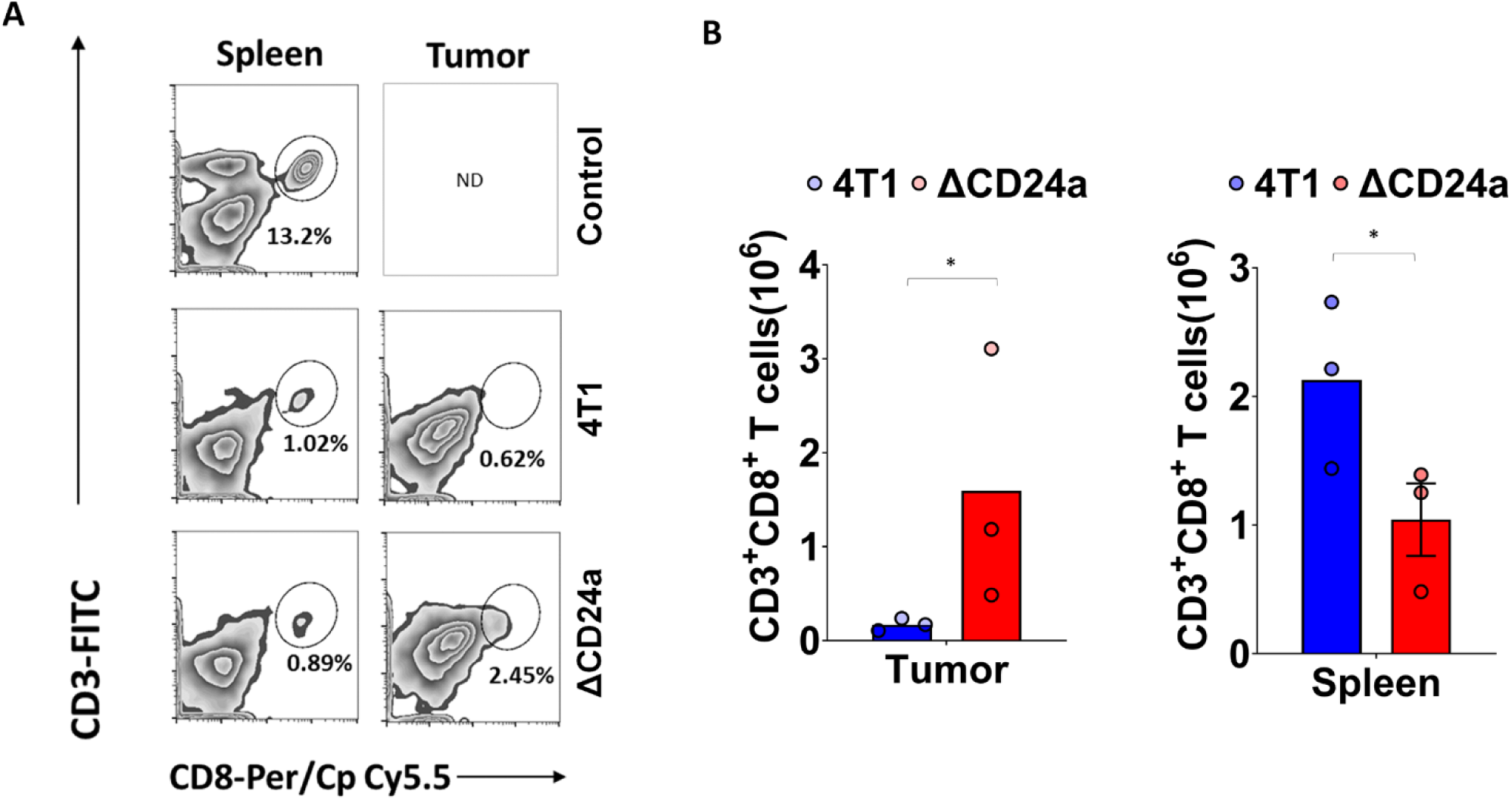
CD24a loss significantly enhances CD8^+^ T cells infiltration into tumors. **A**, Flow cytometry analysis of CD8^+^ T cells in the spleen and tumor (n=3). **B**, The cell numbers of CD8^+^ T cells in the tumor and spleen are presented in histograms. *P<0.05.

### Immunofluorescence (IF) staining confirms the “hotness” of the tumor immune microenvironment of CD24a knockout 4T1 tumors

To further validate the “hotness” of the ΔCD24a 4T1 tumors, we performed IF staining to examine the distribution of tumor-infiltrating macrophages, MDSCs, and cytotoxic CD8^+^ T cells. The results of IF staining revealed a significant increase in the infiltration of F4/80^+^ macrophages into TME of ΔCD24a 4T1 tumors compared to WT tumors (Fig. 5A). Additionally, the infiltration of CD86^+^ M1 macrophages was substantially higher compared to CD206^+^ M2 macrophages (Fig. 5A). The ΔCD24a 4T1 tumors exhibited a substantial reduction of Gr-1^+^ MDSCs infiltration and a significant increase of cytotoxic CD8^+^ T cells infiltration as compared to WT tumors (Fig. 5B). Taken together, the IF results confirmed that ΔCD24a 4T1 tumors exhibited a hot tumor phenotype, characterized by enhanced infiltration of relevant immune cells including CD86^+^ M1 macrophages and cytotoxic CD8^+^ T cells, in contrast to the “cold” phenotype of WT 4T1 tumors.

**Fig. 5.**
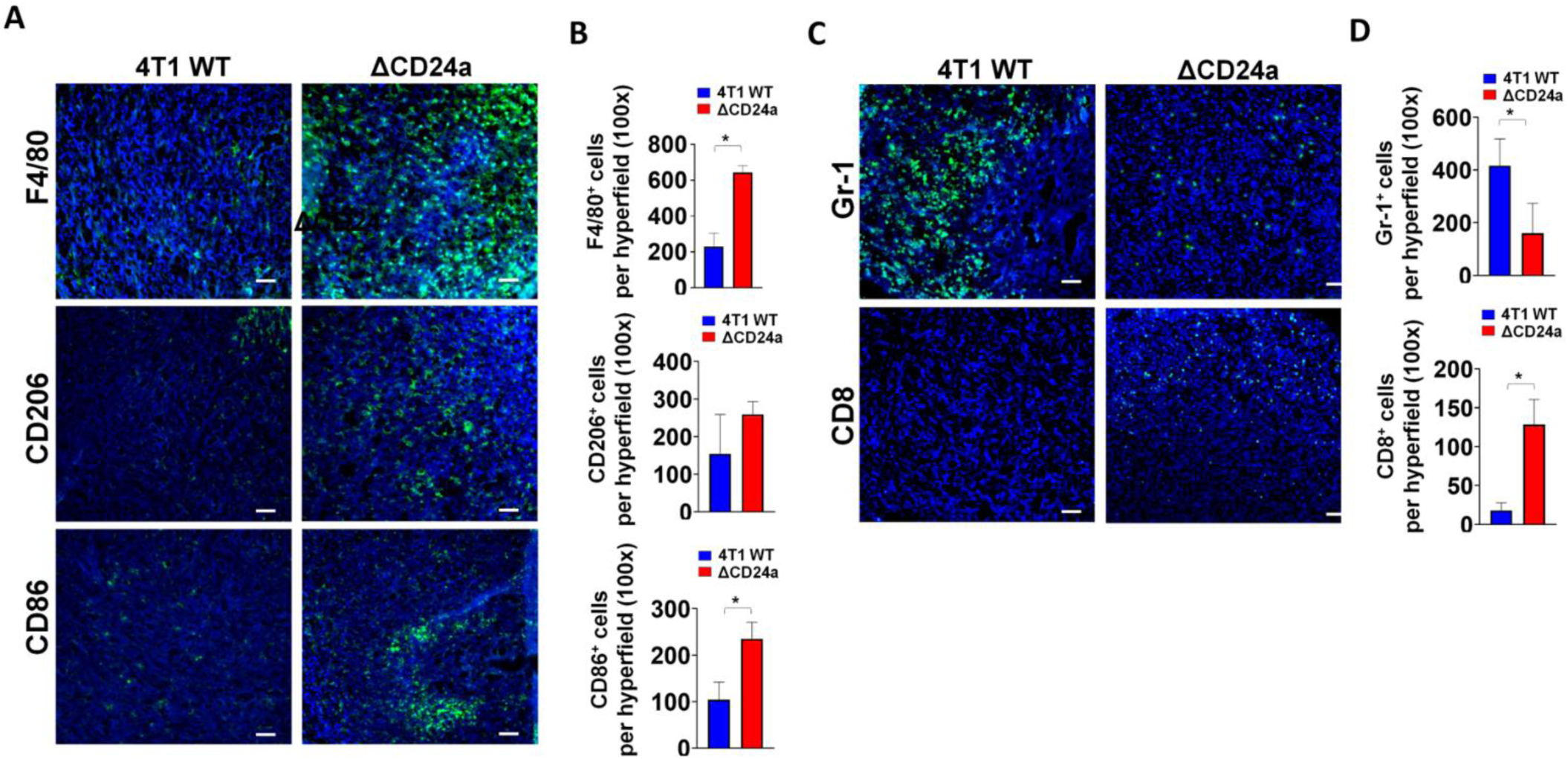
Immunofluorescence staining confirms the transformation of tumor immune microenvironment from a cold tumor to hot tumor state in CD24a knockout 4T1 tumors. **A**, The representative tumor section IF images of F4/80^+^ macrophages, CD206^+^ macrophages and CD86^+^ macrophages. **B**, Quantitative results of tumor section IF images of F4/80^+^ macrophages, CD206^+^ macrophages, and CD86^+^ macrophages. P<0.05**. C**, The representative tumor section IF images of Gr-1^+^ MDSCs and CD8^+^ T cells. **D**, Quantitative results of tumor section IF images of Gr-1^+^ MDSCs and CD8^+^ T cells. P<0.05.

### Three-dimensional (3D) reconstruction of the tumor immune microenvironment (TIME) reveals the detailed information on tumor hotness and coldness

Subsequently, we employed the IF approach and combined it with a tissue clearance technique, utilizing a high-speed confocal microscopy imaging system to create 3D maps of both ΔCD24a 4T1 tumors and 4T1 tumors to further explore the spatial distribution and interactions of tumor-infiltrating immune cells within the tumor microenvironment (TIME) (Fig. 6A). The distribution of Gr-1^+^ MDSCs, F4/80^+^ macrophages, and CD8^+^ T cells within the tumor microenvironment (TME) was assessed. The results of 3D maps of TME revealed that 4T1 tumors exhibited robust infiltration by Gr-1^+^ MDSCs but had very few F4/80^+^ macrophages (Fig. 6B-6D). In contrast, ΔCD24a 4T1 tumors displayed significant infiltration of F4/80^+^ macrophages, instead of Gr-1^+^ MDSCs (Fig. 6B-6D). Moreover, we observed an increased number of tumor-infiltrating CD8^+^ T cells in the ΔCD24a 4T1 tumors (Fig. 6E-6G). These findings further confirm the critical role of CD24a in transforming the TME from a “cold” to a “hot” state.

**Fig 6.**
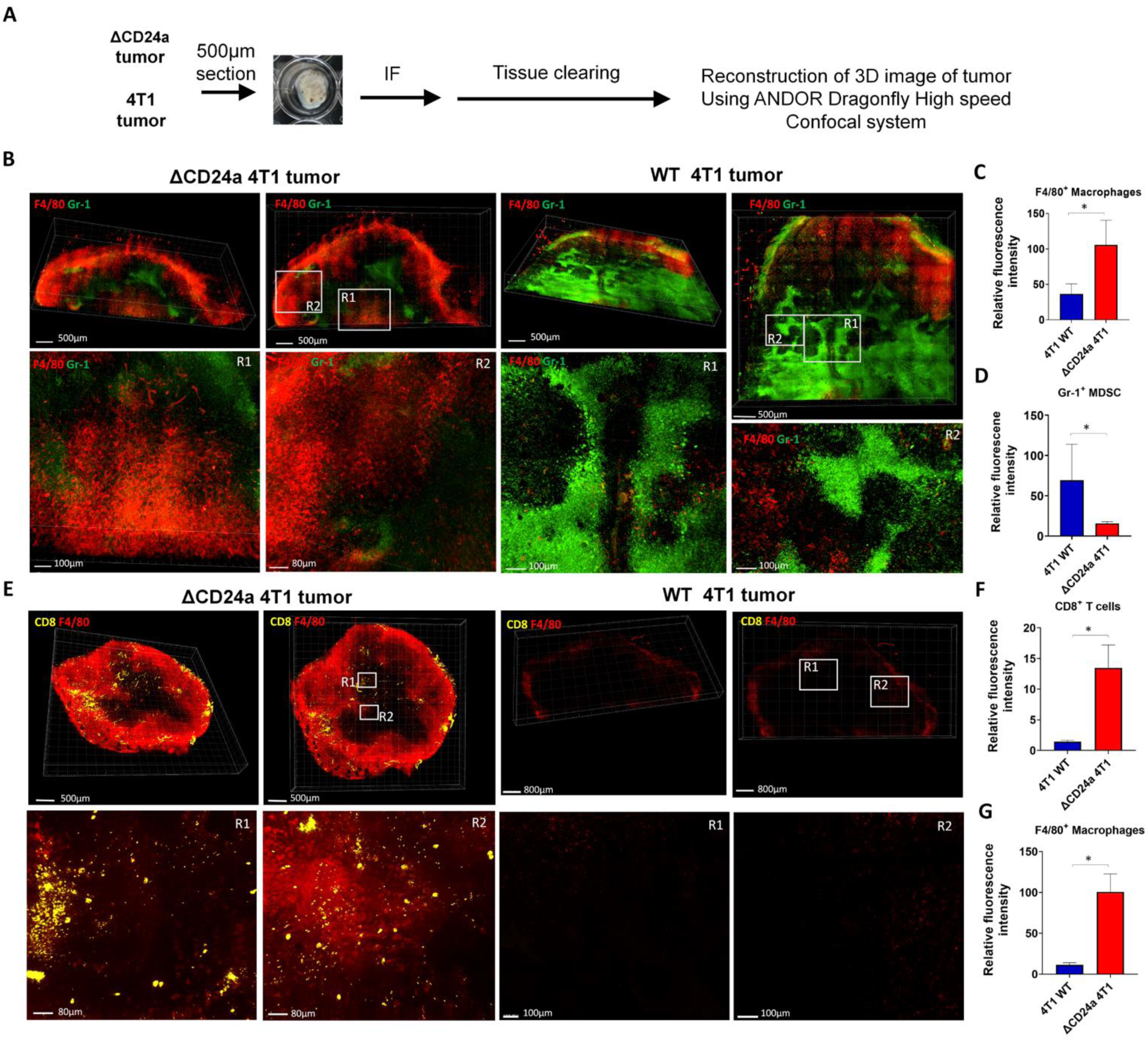
Reconstruction of three-dimensional images validates the hotness of CD24a knockout 4T1 tumors. **A**, Flow chart of reconstruction of 3D images of tumors. **B**, Fluorescence images of F4/80^+^ macrophages and Gr-1^+^ MDSC within 4T1 tumor and ΔCD24a 4T1 tumor. **C-D**, Quantitative results of fluorescence signal of F4/80^+^ macrophages and Gr-1^+^ MDSC within 4T1 tumor and ΔCD24a 4T1 tumor. *P<0.05. **E**, Fluorescence images of F4/80^+^ macrophages and CD8^+^ T cells within 4T1 tumor and ΔCD24a 4T1 tumor. **F-G**, Quantitative results of fluorescence signal of F4/80^+^ macrophages and CD8^+^ T cells within 4T1 tumor and ΔCD24a 4T1 tumor. *P<0.05.

## Discussion

In this study, we investigated the impact of CD24a loss on cancer progression and the immune landscape of the tumor microenvironment (TME) using the mouse carcinoma 4T1, which is a murine model of triple-negative breast cancer (TNBC) and is characterized by poor immunogenic, i.e. an immune cold tumor [20]. Our findings revealed that knocking out CD24a not only significantly hindered the progression of 4T1 tumors but also greatly influenced the immune components of TME. We demonstrated that the absence of CD24a reduced the recruitment of gMDSCs, and significantly increased the infiltration of M1 macrophages, cytotoxic CD8^+^ T cells, and CD49b^+^ NK cells into the TME, transforming the TME from a cold tumor state (poorly immunogenic) to a hot tumor state (highly immunogenic). Consequently, mice bearing ΔCD24a 4T1 tumors exhibited a longer survival time than those bearing 4T1 tumors.

The “hotness” and “coldness” of tumors are defined by many crucial factors including immune contexture of the TME, and cancer cells themselves [21]. TNBC often develops resistance to first-line chemotherapy, and accumulates mutations, leading to the transformation of the TME into a “cold” state, which becomes less responsive to immune checkpoint inhibitors (ICIs) therapy [22]. The molecular heterogeneity of TNBC such as the accumulation of tumor mutation burdens (TMBs) and the activation of pathways like NF-κB, PTEN/PI3K/AKT/mTOR, JAK/STAT, and receptor tyrosine kinases, contributes to chemoresistance and tumor progression [23]. However, targeted therapeutics for these pathways have proven ineffective for TNBC patients [23]. CD24 has recently been identified as a central molecule in tumor immune evasion by interacting with Siglec-10 on macrophages, sending a “don’t eat me” signal that prevents phagocytosis [17]. However, whether CD24 directly contributes to fostering a relative cold tumor immune microenvironment (TIME) is poorly investigated. In this study, we demonstrated that knocking out the murine CD24a gene in 4T1 cells significantly increased the population of tumor-infiltration macrophages in the TME (Fig. 3A), suggesting that the presence of CD24a could effectively inhibit the infiltration of macrophages. Furthermore, our IF results indicated that a substantial increase of CD86^+^ M1 macrophages but not CD206^+^ M2 macrophages was observed in ΔCD24a tumors as compared to 4T1 tumors (Fig. 5A). Additionally, we showed that knocking out CD24a resulted in a substantial increase of CD8^+^ T cell infiltration in the tumors of ΔCD24a 4T1 tumor-bearing mice as compared to that of 4T1 tumor-bearing mice (Fig. 4). The accumulation of cytotoxic T cell infiltration and M1 macrophages in the TME is a hallmark of “hot” tumors [20, 21]. Our results clearly showed that CD24a knockout transforms 4T1 tumors from an immune cold state to a hot state.

In addition, the number of tumor-infiltrating CD11b^+^Ly6G^+^ granulocytic MDSCs (gMDSCs) but not CD11b^+^Ly6C^+^ monocytic MDSCs (mMDSCs) dropped substantially in ΔCD24a 4T1 tumors as compared to 4T1 tumors (Fig. 2A and 2B). Meanwhile, CD24a loss also decreased both gMDSCs and mMDSCs in the spleen (Fig. 2A and 2B). These results suggested that CD24a loss significantly affected the migration of gMDSCs from bone to the TME and to the spleens. Previous studies have shown that both gMDSCs and mMDSCs play significant roles in promoting tumor progression by suppressing cytotoxic T cell activation and proliferation in different ways [24–29]. gMDSCs primarily produce reactive oxygen species (ROS) and nitric oxide (NO) to suppress T cell function [28]. On the other hand, mMDSCs produce immunosuppressive cytokines like TGF-beta and IL-10 and have higher levels of arginase activity, which has been shown to compete arginine as the energy resource with TILs [29, 30]. Our results suggest that knocking out CD24a may negatively affect the expression of chemokines related to the recruitment of gMDSCs in 4T1 cells [31, 32], thereby preventing gMDSC recruitment and expansion. Further investigation is warranted to understand the specific mechanisms by which CD24a regulates chemokine expression and immune cell migration. Additionally, the number of gMDSCs in the TME was approximately thirty times greater than mMDSCs, indicating that gMDSCs play a more significant role in immunosuppression during tumor progression compared to mMDSCs (Fig. 2B and 2C).

In-depth examination of the tumor microenvironment (TME) using 3D reconstruction techniques enabled us to compare immune cell infiltration patterns between 4T1 tumors and ΔCD24a 4T1 tumors. By reconstructing tumor sections with up to 500 μm thickness, this advanced method provides a comprehensive three-dimensional perspective on the spatial distribution and interactions of tumor-infiltrating immune cells. The 3D reconstruction confirmed that 4T1 tumors had substantial infiltration by Gr-1^+^ MDSCs but very few F4/80^+^ macrophages. In contrast, ΔCD24a 4T1 tumors displayed significant infiltration by F4/80^+^ macrophages instead of Gr-1^+^ MDSCs (Fig. 6B-6D). Additionally, ΔCD24a 4T1 tumors showed an increased number of tumor-infiltrating CD8^+^ T cells (Fig. 6E-6G). These findings demonstrate that CD24a plays a crucial role in modulating the TME. Specifically, the absence of CD24a leads to a shift from a suppressive immune environment dominated by MDSCs to an active immune environment with increased macrophage and CD8^+^ T cell infiltration, effectively transforming the TME from “cold” to “hot”. This transformation is significant as it suggests that targeting CD24 could enhance anti-tumor immune responses and improve the efficacy of immunotherapies.

## Conclusion

Our study reveals that CD24a plays a pivotal role in immune evasion and tumor progression in the murine TNBC model. The loss of CD24a significantly impedes tumor growth and transforms the TME from an immune cold to a hot state, characterized by reduced gMDSCs and increased infiltration of macrophages, cytotoxic CD8+ T cells, and NK cells. These findings further support the notion that targeting CD24 could be a promising strategy for converting immune cold tumors into hot tumors, thus improving their responsiveness to current immune checkpoint inhibitors (ICIs). Overall, targeting CD24 presents a novel approach to reprogram the TME, potentially improving the responsiveness of TNBC and other cancers to immunotherapy.

## Availability of data and materials

All data are included in this published article and Additional files.

## Abbreviations

TNBC: triple-negative breast cancer
TME: tumor microenvironment
MDSCs: myeloid-derived suppressor cells
NK cell: natural killer cells
CRISPR: clustered regularly interspaced short palindromic Repeats
ICI: immune checkpoint inhibitor.

## Acknowledgements

We would like to thank Prof. Heng-Hsiung Wu at China Medical University for technical assistance of 3D mapping of tumors.

## Funding

This work was financially supported by the National Science and Technology Council (NSTC) of Taiwan (NSTC 111-2622-B-039-005, NSTC 112-2622-B-039-006), and the Cancer biology and precision therapeutics center and Chinese Medicine Research Center, China Medical University from The Featured Areas Research Center Program within the framework of the Higher Education Sprout Project by the Ministry of Education (MOE) in Taiwan (CBPTC-PROJ-5-2, CMRC-CENTER-0).

## Authors’ contributions

SH-C conceptualized the study, performed experiments, analyzed the data, wrote manuscript, and provided funding; FP-S and HJ-T performed experiments, and analyzed the data; LH-W supervised the study, finalized the manuscript, and provided funding. All authors read and approved the final version of manuscript.

## Ethics declarations

### Ethics approval and consent to participate

All animal experiments received approval from the Animal Use Protocol Committee at China Medical University (CMUIACUC number: 2022-362) and were conducted in compliance with the National Institutes of Health (NIH) guidelines for the Care and Use of Laboratory Animals.

### Consent for publication

Not applicable

### Competing Interests

The authors have stated that they have no competing interests.

## Notes

### Competing Interest Statement

The authors have declared no competing interest.

