## Supplemental files for "CD24a knockout transforms the tumor microenvironment from cold to hot by promoting tumor-killing immune cell infiltration in a murine triple-negative breast cancer model"

**Supplementary Table S1. Antibodies, reagents, and chemicals.**

| Antibodies | Cat. number | manufacturer |
|---|---|---|
| FITC-conjugated anti-CD11b antibody | 101206 | Biolegend, CA, USA |
| PE-Cy7-conjugated anti-Ly6C antibody | 128018 | Biolegend, CA, USA |
| PE-conjugated anti-Ly6G antibody | 164504 | Biolegend, CA, USA |
| FITC-conjugated anti-CD11b antibody | 101206 | Biolegend, CA, USA |
| PerCP/Cy5.5-conjugated F4/80 antibody | 123128 | Biolegend, CA, USA |
| FITC-conjugated anti- $CD49b^+$ antibody | 108905 | Biolegend, CA, USA |
| Accutase | 423201 | Biolegend, CA, USA |
| Ficoll-Paque™ PLUS | 17144003 | Cytiva, CA, USA |
| 0.25% Trpsin-EDTA | 25200056 | ThermoFisher Scientific Inc., CA, USA |
| RapiClear CS solution | RCCS001 | SunJin Lab Co. Taiwan |

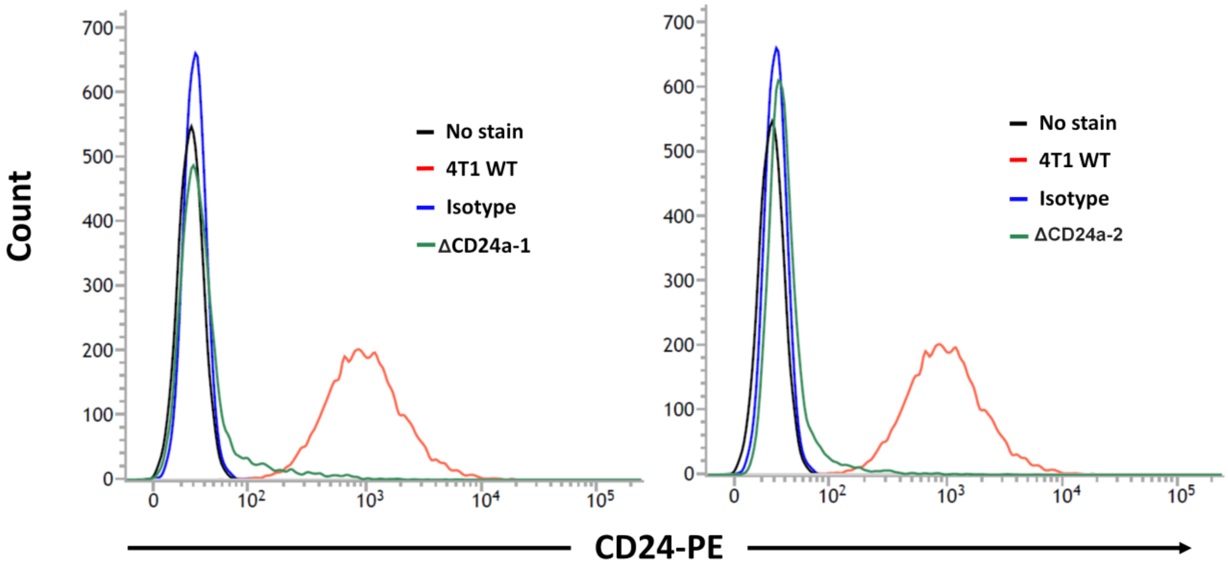


**Supplementary Fig. S1. Flow cytometry analysis of CD24 expression in 4T1 cells and two CD24a knockout 4T1 cells.**

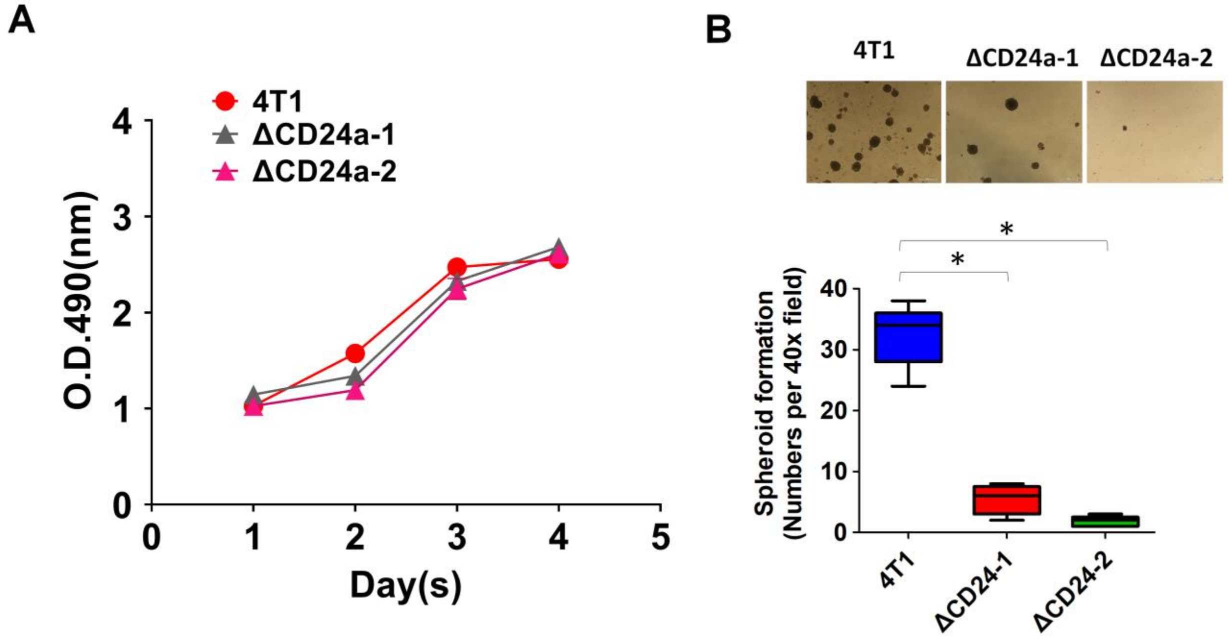


**Supplementary Fig. S2. Evaluation of the effect of CD24a knockout on cell proliferation and spheroid formation. A,** Cell proliferation analysis of 4T1 and CD24a knockout cell using MTS assay. **B,** Tumor spheroid formation analysis of 4T1 and CD24a knockout cell using spheroid culture protocol. * $P<0.01$.
